# reactIDR: Evaluation of the statistical reproducibility of high-throughput structural analyses for a robust RNA reactivity classification

**DOI:** 10.1101/275016

**Authors:** Risa Kawaguchi, Hisanori Kiryu, Junichi Iwakiri, Jun Sese

## Abstract

**Motivation:** Recently, next-generation sequencing techniques have been applied for the detection of RNA secondary structures called high-throughput RNA structural (HTS) analysis, and dozens of different protocols were used to detect comprehensive RNA structures at single-nucleotide resolution. However, the existing computational analyses heavily depend on experimental data generation methodology, which results in many difficulties associated with statistically sound comparisons or combining the results obtained using different HTS methods.

**Results:** Here, we introduced a statistical framework, reactIDR, which is applicable to the experimental data obtained using multiple HTS methodologies, and it classifies the nucleotides into three structural categories, *stem, loop*, and *unmapped*. reactIDR uses the irreproducible discovery rate (IDR) with a hidden Markov model (HMM) to discriminate accurately between the true and spurious signals obtained in the replicated HTS experiments. In reactIDR, IDR and HMM parameters are efficiently optimized by using an expectation-maximization algorithm. Furthermore, if known reference structures are given, a supervised learning can be applicable in a semi-supervised manner. The results of our analyses for real HTS data showed that reactIDR achieved the highest accuracy in the classification problem of stem/loop structures of rRNA using both individual and integrated HTS datasets as well as the best correspondence with the three-dimensional structure. Because reactIDR is the first method to compare HTS datasets obtained from multiple sources in a single unified model, it has a great potential to increase the accuracy of RNA secondary structure prediction at transcriptome-wide level with further experiments performed.

**Availability:** reactIDR is implemented in Python. Source code is publicly available at https://github.com/carushi/reactIDRhttps://github.com/carushi/reactIDR.

**Supplementary information:** Supplementary data are available at online.

## 1 Introduction

RNA secondary structures are known to play diverse roles in the fundamental functions of RNA such as gene expression regulation, translation, and localization, which are mediated by certain structural motifs or regional accessibilities (Borujeni *et al.*, 2013; Nawrocki *et al.*, 2014; Hamilton and Davis, 2011). Many previous studies have succeeded in inferring possible secondary structures or accessibility from both experimental and computational perspectives (Puton *et al.*, 2013). Due to the difficulties in the experimental determination of RNA secondary structure prior to the development of the next-generation sequencing, a number of computational methods was developed to predict the secondary structure from sequences or sequence alignment data (Zuker *et al.*, 1989) These methods were shown to be highly accurate in the small RNA analyses, but the time and computer memory requirements remain too high to be applicable at genome-wide level (Bernhart *et al.*, 2006; Kawaguchi and Kiryu, 2016).

To discover a comprehensive landscape of RNA secondary structure, novel types of high-throughput experimental methods, such as PARS (Kertesz *et al.*, 2010) and icSHAPE (Spitale *et al.*, 2015), have been developed using short-read next-generation sequencers, referred to as high-throughput RNA structural (HTS) analysis. Those experiments do not predict RNA secondary structures directly, but the scores obtained, termed reactivity scores, can provide us the information about molecular structures. The reactivity scores can provide *in silico* methods with information to guide secondary structure prediction (Deigan *et al.*, 2009), as well as directly indicate the propensity of structural accessibility (Taliaferro *et al.*, 2016). The reactivity scores are obtained by the distribution of chemical or enzymatic footprints of RNA structural elements obtained from the HTS data. As summarized in Figure 1(A), each experimental method detects different types of structural footprints to determine single- or double-strandedness of each nucleotide. For instance, PARS method employs RNase V1 and S1, which leads to a structure-specific cleavage at the stem (base-paired) and loop (unpaired) regions in the RNA molecules, respectively (Kertesz *et al.*, 2010). Conversely, using icSHAPE (Spitale *et al.*, 2015), one of the SHAPE-Seq(Lucks *et al.*, 2011)-like methods, base modifications are introduced in the loop regions by NMIA-*N*_3_ treatment, and this modification is then detected according to a frequent drop-off at the spot during reverse transcription (RT) with efficient streptavidin pull-down purification. With the exception of a novel type of methods, in which mutational profiling information is comprehended as a modification clue (Siegfried *et al.*, 2014; Zubradt *et al.*, 2017), most of the existing HTS approaches are classified into the three types presented in Figure 1(A) (Lorenz *et al.*, 2016).

**Figure 1:**
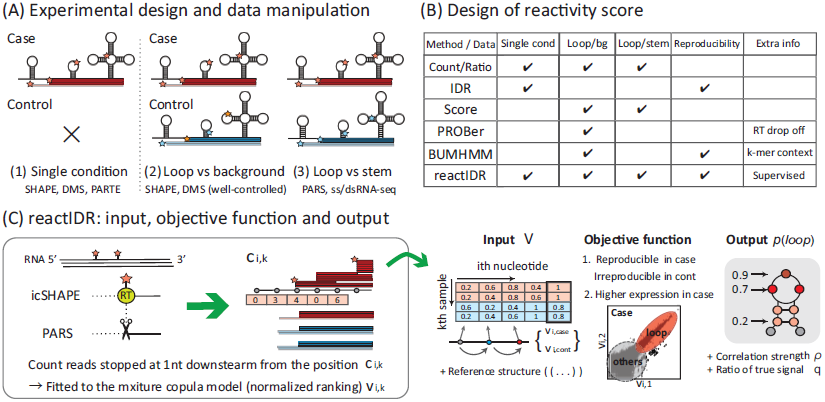
Schematic illustration of HTS analyses and reactIDR. (A) Major types of HTS experimental designs analyzed in this study. Each approach can be classified into the presented types based on the number of sample conditions and their detection targets. (B) Applicability of reactivity-scoring methods to three types of HTS designs (*i.e.*, whether the method is intended or not), in which *Score* indicates each threshold-based scoring method used for studying the icSHAPE (Flynn *et al.*, 2016) and PARS (Wan *et al.*, 2014) experiments. (C) Overview of reactIDR. reactIDR receives the input data consisting of 5′-end read depth repeatedly measured in a single or two conditions. The input data, after trimming of the very end of transcripts, is converted into the scaled ranking data to adapt the joint cumulative probability distribution of copula model. Afterward, a posterior loop probability distribution at each site can be computed by using reactIDR as an index for reactivity.

While these HTS methods rely on different approaches, they suffer from a common problem of experimental noise during the sequencing process. Although the raw footprint levels detected using HTS analyses reflect the strength of the reaction, they simultaneously contain the spurious signals attributed to the reasons other than experimental modifications (*e.g.*, stochastic drop-off during RT, endogenous base modification or cleavage, and sequencing error). To reduce the misdetection of false positives, a naïve way is to compute reactivity scores using the (relative) differences or ratios of the read levels between different experimental conditions, such as case/control (*i.e.*, chemically or enzymatically treated samples *vs.* untreated ones) or V1/S1treated, and afterward, an arbitrary minimum threshold can be manually set to remove false positives (Rouskin *et al.*, 2014; Wan *et al.*, 2014). It is, however, very labor intensive as well as not statistically justified to set a single threshold for each dataset independently, as it may lead to overlooking significant findings or an excessive number of false positives. In more sophisticated methods, probabilistic models, such as Spats (Aviran *et al.*, 2011), ProbRNA (Hu *et al.*, 2013) and PROBer (Li *et al.*, 2017), were also introduced, in which the detection of the regions with different read depth between case/control samples is most heavily weighed. Despite achieving background noise reduction by using the probabilistic models, these methods have issues related with the utilization of reproducibility information between the replicates, which general high-throughput sequencing analyses have (Robinson *et al.*, 2010; Li *et al.*, 2011).

Consistency between replicates generally indicates the quality of the experiments beyond stochastic noise, particularly for the sequencing-based experiments. Therefore, in almost all sequencing approaches, the reliability of their conclusions is evaluated based on the consistency and correlation between replicates. Using these findings, it is possible to enhance the accuracy of downstream analyses, such as in *in silico* structure prediction or comparison between multiple datasets. Several statistical methods have been developed to estimate the reliability of HTS analysis reactivity using the replicate information, such as Mod-seeker (Talkish *et al.*, 2014) and BUMHMM (Selega *et al.*, 2016). Mod-seeker is based on the ratio of footprint counts between case and control and does not utilize a sophisticated probabilistic model. On the other hand, BUMHMM closely depends on a certain statistical model based on the empirical distribution fitted to the target experiments, such as SHAPE-Seq and DMS-Seq. Due to the diversity of HTS methodologies, the distribution of structural footprints is known to be highly heterogeneous (Supplementary Figures 1 and 2), so there is no guarantee when applying the model to other HTS methodologies.

**Figure 2:**
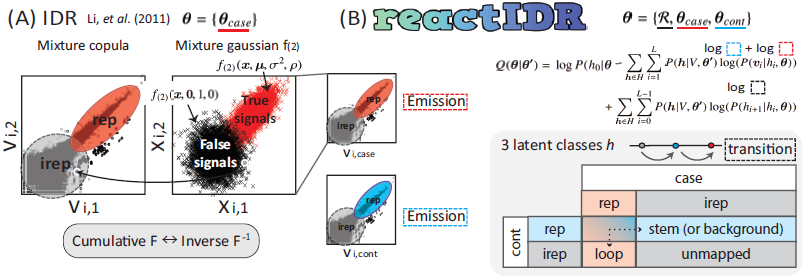
The comparison between the irreproducible discovery rate (IDR) and reactIDR model. The IDR is a method for the classification of signals into true (high-coverage and reproducible) or false (low-coverage and irreproducible) signals, based on the Gaussian mixture model, by associating the observation with the cumulative joint probability distribution and pseudo-value *x*. In reactIDR, IDR is combined with the hidden Markov model of the three latent classes (*loop, stem*, and *unmapped*) to handle case/control comparisons with the Gaussian mixture copulas for each condition simultaneously, in which each parameter is optimized to the data and reference structure by expectation-maximization algorithm.

The importance of a comprehensive HTS comparison was reported previously, in a study showing that the results of each HTS approach are less correlated than those from the same approach, and potentially contain largely non-overlapping conformational information (Sexton *et al.*, 2017). Therefore, a fair comparison of multiple experiments requires unified statistical reliability, whereby the results from different experiments can be merged and compared, such as *in vivo* and *in vitro* results, or results obtained using different approaches (Choudhary *et al.*, 2017).

To overcome HTS dataset heterogeneity, one promising approach is combining a probabilistic model and supervised learning. A reliable structural information has been gathered in previous computational and experimental analyses (Nawrocki *et al.*, 2014) and many previous studies have applied supervised learning to *in silico* structure prediction to estimate the optimal model parameters and improve the accuracy of structural prediction. However, to the best of our knowledge, there is no method to evaluate the unified reliability of reactivity scores obtained from the HTS approach in a supervised learning manner (Figure 1(B)). As such, there is room to improve in computational methods for HTS analysis, especially by allowing the handling of multiple HTS experiments simultaneously, for comprehensive understanding of RNA secondary structure.

Here, we have developed a novel method, reactIDR, to determine true reactivity signals using the replicated HTS data by calculating a statistically valid reliability score. To evaluate the reliability in a way applicable to a general HTS dataset, we extend a statistical method for chromatin immunoprecipitation (ChIP)-Seq peak detection, named irreproducible discovery rate (IDR) (Li *et al.*, 2011). Although the existing IDR-based methods regard all observed points as independent, similar to the ChIP-Seq peak signals, sequence read numbers covered on the consecutive nucleotides depend on each other in the HTS datasets. Hence, for the application of IDR to HTS, we have applied a hidden Markov model (HMM) with the emission probability of IDR, in which the loop and stem regions are automatically segmented by a maximum posterior estimate. reactIDR is the first framework for supervised learning with known secondary structures obtained from HTS datasets as well as applying parameter optimization based on the expectation-maximization (EM) algorithm to avoid any arbitrary or time-consuming parameter hand-tuning step,

## 2 Materials and Methods

### 2.1 reactIDR overview

Our novel pipeline, reactIDR, was designed to explore optimal stem/loop classification from data obtained in various HTS analyses, according to IDR criteria (Li *et al.*, 2011). A schematic workflow of reactIDR is presented in Figure 1(C) (and Supplementary Figure 3). We aimed to develop reactIDR so that it can be applied for solving HTS problems as follows:

**Figure 3:**
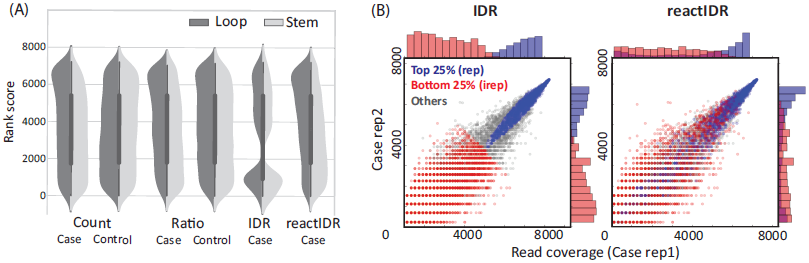
Distribution of raw read count and reactivity scores of reactIDR. (A) Violin plots, showing the ascending rank orders of the four indices of chemical footprint enrichment that can be computed for case and control samples separately: *Count*, raw read count; *Ratio*, read count normalized by the reads passing through the position; irrepro- ducible discovery rate (IDR); and reactIDR. With the increase in the rank of the reactivity score obtained from the case samples, the likelihood of the nucleotide being in the loop structure increases. (B) Read count distribution of reproducible and irreproducible groups, classified into top 25 % and bottom 25 % rankings, based on the IDR (left) and reactIDR reactivity (right) criteria.

- **Input** - Assuming that *K* is the number of samples in an HTS experiment, and sequencing reads were obtained from two *K/*2 samples, in two different conditions (and an optional reference structures). Each read is mapped against a reference sequence, and the indicator of reactivity at each base is measured by counting the reads that start at the subsequent (3′) base.
- **Output** – The outputs represent the probabilities that each nucleotide is the latent class *loop*.
- **Objective function** – To provide reliable reactivity evaluation and accurate structure prediction, the outputs should be optimized so that they satisfy the following conditions:(1)the sequential loop regions of RNAs provide consistently enriched read counts in the case condition, in contrast to those in the control conditions, while stem motifs should be enriched in the control obtained from the PARS dataset, and (2) the true signals of structural footprints should be consistent among the replicates with expression increased as much as possible.

In the HTS approach, the positions in stem (base paired) or loop (accessible) structures can be detected by the enrichment of chemical or enzymatic footprints, since their reactivity is highly affected by the existence of base pairs. However, to remove the considerable levels of false positives derived from the random occurrences of fragmentation or endogenous modification, reactIDR further considers the ratio of read count enrichment between two-conditional samples. As shown in Figure 1, *case* (*control*) samples were specifically defined as the first (last) *K/*2 samples with the sequencing reads stochastically truncated at one base downstream from the loop (stem or non-specific). In reactIDR, a latent class *loop, stem*, and *unmapped* is expected to be assigned to the region corresponding to case-derived, control-derived, and no sufficient reads are obtained, respectively. Here, icSHAPE case and control samples were regent-treated and untreated, respectively, while PARS samples were treated with S1 and V1, respectively, and there, the control samples were not untreated samples. Using reactIDR, we can compute a posterior probability distribution for each latent class at each site as an index of the reactivity.

### 2.2 IDR

Our novel algorithm reactIDR extends the idea of IDR (Li *et al.*, 2011) criteria to obtain reproducible and irreproducible classification, a typical problem of high-throughput sequencing beyond the fundamental problem of irreproducible read sampling, hence we first introduce the model for IDR estimation. Here, as an input of IDR and reactIDR, let us consider read-count data obtained for a single transcript or concatenated multiple transcripts with the total length of *L* nucleotides. As the read count data is initially converted into ranking data across all positions in each experiment, this ranking data in ascending order is scaled to the range of [0, 1]. This normalized data is hypothesized to correspond to the cumulative probability distribution in the copula model, necessarily being within that range. After scaling, a vector of the ranks of read counts{at the *i*th position is defined as ***v***_*i*_:={{*v*_*i,*1_, *…, v*_*i,K*_}} for IDR, and*vi*:= {*v*_*i,case*_, *v*_*i,cont*_} ={*v*_*i,*1_, *…, v*_*i,K/*2_}, {*v*_*i,K/*2+1_ *… v*_*i,K*_} for reactIDR. The overall input data of IDR and reactIDR is *V* ={ ***v***_1_, *…,* ***v***_*L*_.}

The basis of IDR is using the mixture copula model to discriminate one reproducible (represented by *rep*) distribution from the irreproducible (*irep*) distributions, recognizing them as true and false signals, respectively. True signals obtained from the protein binding sites in ChIP-Seq are supposed to exhibit reproducible enrichment with an increase in the read-count depth and a high correlation between the replicates. Based on this hypothesis, the signals categorized into *rep* are considered to contain many more true signals than the *irep* group.

For IDR estimations, a mixture copula is assumed, in which each copula can explain the distribution of true reproducible signals, except for a single copula corresponding to the false signals that are poorly or not correlated between replicates. For simplicity, we considered an example case of the mixture of two Gaussian copulas in two dimensions hereafter, which can handle a case of two experimental replicates. When the *i*th signal is produced by true (false) signals, represented as *r*_*i*_ = *true*(*false*), the distribution of a random variable *x*_*i*_ = (*x*_*i,*1_, *x*_*i,*2_) corresponding to the *i*th signal is as follows:

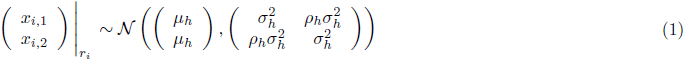

where *μ*_*h*_, *σ*_*h*_, and *ρ*_*h*_ represent mean, variance, and correlation, respectively, with the indicator *h* for each state of *r*_*i*_ (*h* = 1 when *r*_*i*_ = *true* and *h* = 0 when *r*_*i*_ = *false*). The distribution of true *rep* signals is assumed to contain a positive mean and correlation (*μ*_1_ *>* 0 and 0 *< ρ*_1_ ≤1) while spurious *irep* signals should be located around the low coverage regions and show increased variance (*μ*_0_ = 0,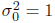, and *ρ*_0_ = 0). These parameters can be fitted, not to the actual read counts, but to the normalized ranks of read coverage for ChIP-Seq data ***v***, such that those ranks are derived from the cumulative marginal probability distribution function of *x*_*i,k*_ (*k* = 1, 2), as shown below:

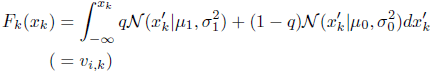

where *q* represents the ratio of true and reproducible signals in the samples. In this way, a probability of each peak, derived from the irreproducible signal or IDR, can be estimated.

### 2.3 reactIDR: A novel algorithm for the extraction of reproducible signals based on the IDR and HMM

While IDR is a powerful method for the evaluation of the reliability of various joint distributions of ChIP-Seq peaks, HTS data used for RNA stem/loop detection must allow comparisons between multiple conditions considering the dependency between the consecutive nucleotides. The application of IDR to HTS, therefore, requires further specific extensions for the more complicated situations. To this end, we developed reactIDR, a novel method for the extraction of true reactivity signals from replicated HTS data, based on the determination of statistical reliability score according to IDR and HMM.

For *L* as a total length of transcripts and a vector of scaled rankings at the *i*th position, ***v***_***i***_ can be given. In reactIDR, a latent variable at the *i*th position *h*_*i*_ can belong to any element of {*loop, stem*, and *unmapped*} corresponding to each status associated with HTS data and RNA secondary structures. The enrichment of ***v***_***i***_ is thought to be most likely observed at the regions that belong to the *loop* class in the case samples and *stem* class in the control samples, while we did not expect any specific enrichment in the case of *unmapped* classes.

In reactIDR, the path of all latent variables are represented by ***h*** ={ *h*_0_, *…, h*_*L*_}, and the likelihoods of *V* and ***h*** can be obtained as the product of emission and transition probabilities, as formulated below:

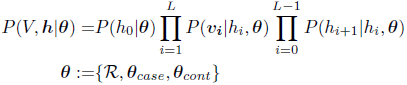

where 𝓡 is a transition matrix between each latent class pair, and *θ*_*case*_ and *θ*_*cont*_ corresponds to the set of all of parameters (*i.e.*, ***μ***, *σ*^2^, and *ρ*) in the copula model for case and control samples, respectively (see Supplementary Figures 4 and 5). Due to the difficulties in the mapping of reads into the edge of transcripts, *h*_0_ and *h*_*L*_ are always assigned to the *unmapped* class (*P* (*h*_0_|***θ***) = 1and *P* (*h*_*L*_|***θ***) = 1). *P* (*h*_*i*+1_|*h*_*i*_, ***θ***) represents the transition probability between *h*_*i*_ and *h*_*i*+1_ in the HMM, represented by *R*, which is further optimized by the expectation of the transition between *h*_*i*_ and *h*_*i*+1_ at each step of the EM algorithm iteration. *P* (***v***_*i*_ |*h*_*i*_, ***θ***) is an emission probability, which we defined as follows:

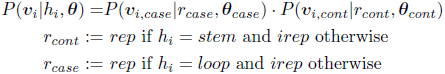

where ***v***_*case*_ and ***v***_*cont*_ are expected to show the enrichment of loop-like and stem-like (or background noise) regions, respectively. These probabilities for *rep* and *irep* signals are obtained by the mixture Gaussian copula as they are in IDR.

A joint distribution *f*_(2)_, cumulative marginal distribution *F*_*k*_, and function of the vectorized inverse of *F*_*k*_ *F* ^*-*^1^^ can be defined as follows:

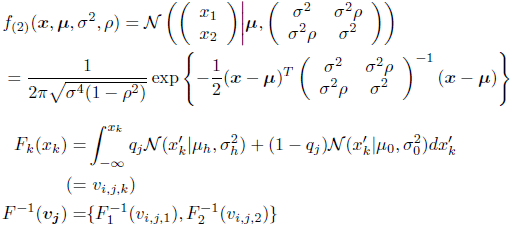

where *h* = 1 (2) when *j* = *case* (*cont*), and *q*_*j*_ represents the ratio of true signals in *j* samples. Afterward, the emission probability based on the mixture Gaussian copula consisting of *rep* and *irep* distributions with parameters ***θ***_*case*_ or ***θ***_*cont*_ can be formulated as follows:

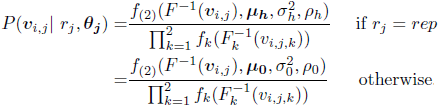

where *μ*_*k*_ *>* 0 and 0 *< ρ*_*k*_ ≤1 while *μ*_0_ = 0, 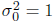, and *ρ*_0_ = 0, the same as those in Eq.1 (detailed in the Supplementary Materials). In this way, using reactIDR, we can computes a posterior probability distribution for each latent class at each site as an index of the reactivity and evaluation of its reliability.

**Figure 4:**
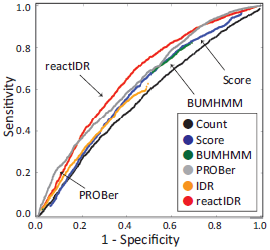
Receiver operating characteristic curve, showing rRNA structure prediction based on the reactivity scores obtained using six scoring methods, for the *in vivo* icSHAPE dataset.

**Figure 5:**
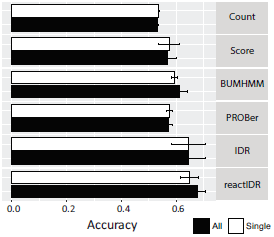
Prediction accuracy of stem/loop structures based on the reactivity scores computed from individual or combined datasets (the icSHAPE *in vivo* and *in vitro* and PARS datasets) by support vector machine, with indeterminable scores filtered out. Average and variance of accuracy were computed after 10-fold cross-validation. The prediction from the single dataset utilized the reactivity scores from either the *in vivo* icSHAPE dataset, *in vitro* icSHAPE dataset, or the PARS V1 dataset, and the results of the maximum mean accuracy are presented here.

### 2.4 Optimization of the reactIDR parameters using the EM algorithm and supervised learning

Using the EM algorithm, each parameter can be iteratively optimized to maximize *Q* in reactIDR defined as follows:

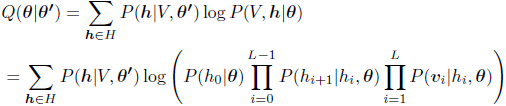

where *H* represents the set of all possible paths of latent variables ***h***, and *θ*′ represents the set of parameters obtained in the previous iteration. Each optimization step is performed by the *fmin l bfgs b* function in SciPy library. Furthermore, reactIDR can also incorporate supervised learning at this step, by limiting *H* so that it is consistent with the reference structure.

In this study, for efficient parameter estimation, reactIDR first accepted positive and negative (or complementary sequences expected not to be transcribed) data associated with 18S rRNA. Afterward, trained parameters were used as the initial set of parameters in the re-fitting process for each rRNA sequence of the test set. The details of the optimization process, such as the derivative of *Q* for each parameter, are described in the Supplementary Materials.

### 2.5 Datasets used and the evaluation of reactivity-based classification

In this study, datasets generated by using two HTS methods, icSHAPE and PARS, were used to validate the accuracy of reactIDR. A PARS-score whole-transcriptome dataset was obtained for the native deproteinized transcriptome of GM12878 cells *in vitro* (GEO accession number,GSE50676) (Wan *et al.*, 2014). Here, we used the normalized read counts of two replicates treated with nucleases S1 and V1 (hereafter referred to as S1 and V1, respectively). Analyses were also conducted using the icSHAPE dataset obtained for the HEK293T cells (GEO accession number, GSE74353), which contains sequencing reads obtained for three conditions with two replicates: dimethyl sulfoxide (DMSO)-, *in vitro* NAI-N3-, and *in vivo* NAI-N3-treated cells (Lu *et al.*, 2016). Computation of the original PARS and icSHAPE scores was implemented following the instructions provided in Wan et al. (Wan *et al.*, 2014) and Flynn et al. (Flynn *et al.*, 2016). The characteristics of these datasets can be found in the Supplementary Materials (Supplementary Figures 6 and 7).

**Figure 6:**
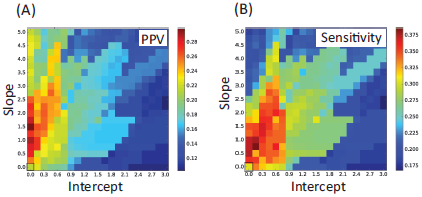
Positive predictive value (PPV) and sensitivity of *in silico* structure prediction with and without reactivity assisting for the test set of rRNAs. The rectangles located at the bottom left corner indicate PPV and sensitivity of fully *in silico* prediction, which is 0.214 and 0.305, respectively. The best PPV and sensitivity pair, selected to maximize their sum was (0.290, 0.373) for reactIDR and (0.249, 0.369) for PROBer, respectively.

**Figure 7:**
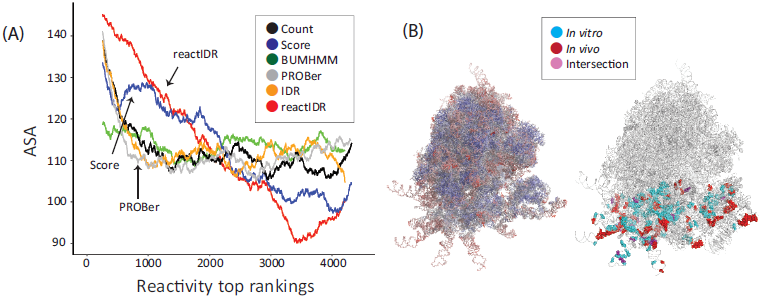
Correlation between RNA reactivity and 3D accessibility. (A) Relation between the accessible surface area (ASA) and reactivity scores computed by six types of scoring methods. y-axis, averaged ASAs within each 500-nucleotide sliding window; x-axis, a ranking of each nucleotide based on the reactivity scores, in which ties were randomly broken. The reactivity scores of reactIDR from the *in vivo* icSHAPE dataset presented the high- est correlation with ASAs across the six scoring methods. (B) 3D structure of ribosomes visualized by PyMOL (Schrödinger, LLC, 2015). The ribosome shown on the left changes its color from red to blue according to the ASAs of each nucleotide (high to low). The highlighted regions on the right indicate the top 7 % reactive sites of 18S rRNA from the dataset of *in vivo* (red), *in vitro* (blue), and their intersection (purple).

We aligned sequencing reads from HTS datasets with bowtie2 (Langmead and Salzberg, 2012) using a reference sequence of human transcriptome and rRNA. The human transcriptome reference consisted of UCSC Refseq sequences (as of October 7, 2016) and GENCODE transcript sequences (v12), with the reference database constructed as described by Wan et al. (Wan *et al.*, 2014). To construct the reference set of the rRNA sequences, a human ribosomal repeating unit (NT 167214.1) was extracted from the NCBI database, and a 5S rRNA sequence (ENST00000364451) was additionally downloaded from the Ensembl database. The process of read counting from the sequencing read data already aligned to the reference sequence was performed by using the Docker image, published with reactIDR. As a rRNA structure reference, cryo-EM-based ribosomal structure (PDB ID: 4v6x) was aligned to our reference sequence(Anger *et al.*, 2013). To obtain base-to-base correspondence between the reference sequences and the structure, the 5S, 5.8S, 18S, and 28S rRNA reference sequences were aligned to each sequence within the structure dataset, and all bases successfully aligned to the reference were used for the evaluation of classification. Of the four rRNAs in the cryo-EM ribosomal structure, 18S rRNA reference structure was used as the training set for the supervised learning of reactIDR, with the negative data for the minus strand of 18S rRNA. The annotations of secondary structure and base pairings were obtained for this structure using RNAview (Yang *et al.*, 2003) with the RNApdbee interface (Antczak *et al.*, 2014). The accessible surface area (ASA) of the ribosome structure was calculated for each nucleotide using NACCESS with default parameter settings (Hubbard and Thornton, 1993).

To evaluate the accuracy of structure classification based on reactivity, we determined a receiver operating characteristic (ROC) curve. *TP, TN, FP*, and *FN* represented the number of true positives, true negatives, false positives, and false negatives. In the ROC curve, the y-axis corresponds to the true positive rates ((*TP*)*/*(*TP* + *FN*)), while the x-axis corresponds to the false positive rate ((*FP*)*/*(*FP* + *TN*)). P-values were computed in order to compare the area under curve (AUROC) of reactIDR and other scoring methods by using the Proc library in R with a bootstrap method with 100 repetitions. We measure the accuracy of *in silico* structure prediction with or without the reactivity scores, positive predictive value (PPV) (*TP*)*/*(*TP* + *FP*), sensitivity (identical to the true positive rates), and Matthews correlation coefficient 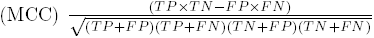for each possible base pair within the transcript were determined. At that point, pseudo-knot and inter-transcript base pairs were converted into the loop structures in the reference dataset. The accuracy of the structure predictions based on multiple HTS datasets was also investigated using support vector machine (SVM) via the interface of the scikit-learn library. For *in silico* structure prediction assisted by reactivity scores, RNAfold from the Vienna RNA package v2.4.0 (Lorenz *et al.*, 2011) was applied with the default parameters.

## 3 Results and Discussion

### 3.1 Characteristics of IDR-based structure classification with or without HMM

We investigated the improvements on combining HMM with IDR as criteria for stem/loop classification in reactIDR. To compare the read coverage distribution of HTS analysis between the case/control conditions and stem/loop positions, *in vivo* icSHAPE data was aligned to the reference sequence of rRNAs and read count (*Count*) was obtained for duplicated case and control samples. The reactivity scores based on *Ratio* (normalized read count ratio versus the number of reads passing through the position), IDR, and reactIDR were also computed from the distribution of *Count* scores. The rank orders from the indices of IDR and reactIDR and averaged indices for *Count* and *Ratio* were then classified into stem/loop locus to examine the existence of read count enrichment amplified by chemical modifications. The locations of loop and stem structures were predicted for four human rRNAs (*i.e.*, 5S, 5.8S, 18S, and 28S) from the 3D structure of ribosome determined by cryo-EM using RNAview (Yang *et al.*, 2003) (detailed in Materials and Methods).

**Table 1:** Area under the ROC curve (AUROC), showing stem/loop classification of reactivity scoring methods.

| Method | icSHAPE |  | PARS |  |
| --- | --- | --- | --- | --- |
|  | *in vivo* | *in vitro* | V1(cont) | S1(case) |
| Count | *0.555 | *0.514 | * 0.702 | * 0.349 |
| Score | *0.600 | *0.564 | † * 0.574 | † 0.574 |
| BUMHMM | *0.592 | *0.526 | † * (0.634) | † (0.634) |
| PROBer | 0.645 | 0.607 | † * (0.638) | † (0.639) |
| IDR | *0.563 | *0.518 | * 0.664 | * 0.375 |
| reactIDR | **0.665** | **0.626** | **0.715** | 0.420 |

\* represents that the area under the ROC curve (AUROC) is significantly lower than that of reactIDR with $p$ -value $< 0.05$ for the comparison after Bonferroni correction. † indicates that the reactivity used for the evaluation is common for the case (loop-enriched) and control (stem-enriched) conditions. The brackets indicates the AUROCs obtained by the methods not originally designed for PARS experiments.

Figure 3 (A) shows the distributions of the rank orders from the indices of chemical footprint enrichment for case and control samples individually. The distributions of *Count* and *Ratio* rank scores of the treated samples were shown to contain a greater number of high-ranking scores to a similar extent each other, compared to those of the control samples. The distribution of IDR rank scores demonstrated two separated clusters associated with reproducibility, due to the characteristics of two Gaussian mixture copula applied in the IDR method (Figure 3(B)). While *Count, Ratio*, and IDR only evaluated the read count enrichment of the samples belonging to one condition, the reactivity scores of reactIDR were estimated based on the case/control ratios and 18S rRNA reference structure. Consequently, reactIDR generated a more distinguishable distribution of higher-ranked scores at the loop regions, compared with that at the stem regions. Since the top rankings for reactIDR tend to contain the nucleotides with not only high read coverage (Figure 3(B)), using reactIDR, we were able to fit more complicated enrichment patterns considering the read coverage of surrounding nucleotides.

Moreover, even for the noisy genome-wide analysis of the PARS data, IDR-based classification was also successful in associating the reproducibility of read enrichment and the strength of stem probability obtained *in silico* (Kawaguchi and Kiryu, 2016) (shown in Supplementary Figure 8). Taken together, using reactIDR case and control read enrichment criteria, we were able to infer the accessibility of each nucleotide with higher precision than that possible when using a raw read count, by employing the data on the reproducibility and reference structure.

### 3.2 Comparison of reactIDR structure classification accuracy for 2D accessibility

To evaluate the accuracy and robustness of reactIDR for the different HTS methodologies, reactivity-based stem/loop classification using multiple HTS rRNAs datasets was performed. For a fair comparison, only 18S rRNA data were subjected to parameter optimization using reactIDR and other rRNAs were subjected to comparison. In addition to the reactivity indices used in the previous analyses, three more types of reactivity scoring methods were computed for each dataset: *Score* used in the original HTS studies, BUMHMM (Selega *et al.*, 2016), and PROBer (Li *et al.*, 2017). Afterward, the AUROC was computed for rRNA structure classification while the structure status expected to be enriched (*i.e.*, stem or loop) was set to positive.

AUROCs for each classification obtained using icSHAPE and PARS datasets are shown in Table 3. reactIDR was observed to generate the highest AUROC for the icSHAPE datasets and the null hypotheses considering the difference in the AUROC obtained by reactIDR and PROBer were rejected for the icSHAPE *in vivo* and *in vitro* datasets (Table 3), indicating the prediction accuracy of reactIDR and PROBer is statistically comparable for the icSHAPE datasets. This consistency was also confirmed for different choices of the training set as well (Supplementary Table 2 and 3). In Figure 5, ROC curves of six reactivity-based predictions for the *in vivo* icSHAPE dataset are presented (see also Supplementary Figures 9 and 10). Using reactIDR, we achieved the highest accuracy for the medium portions (0.1 − 0.6) among the overall range of 1-specificity values, shown along the x-axis. For the stem (V1) prediction from the PARS datasets, reactIDR also generated the highest AUROC among all reactivity scoring methods. On the other hand, the prediction accuracy of all available methods was lower than that randomly obtained in the PARS S1 dataset unless the common reactivity scores were computed for stem/loop prediction (represented by †), as the enrichment of the read count was not observed at loop regions in the PARS S1 dataset (Supplementary Figure 11). Therefore, it can be concluded that reactIDR can be used for the classification of the stem/loop structures, maintaining the sensitivity and specificity, which are enough comparable or higher than those of *Score* and PROBer.

**Table 2:** Correlations between reactivity score and 2D or 3D accessibility indices.

| Method | vs 3D |  | vs 2D |  |
| --- | --- | --- | --- | --- |
|  | *in vivo* | *in vitro* | *in vivo* | *in vitro* |
| Count | *0.099 | *-0.109 | *0.184 | *0.132 |
| Score | *0.207 | -0.033 | *0.207 | *0.175 |
| BUMHMM | 0.021 | *-0.155 | *0.194 | *0.111 |
| PROBer | *0.096 | *-0.092 | *0.234 | **\*0.205** |
| IDR | *0.105 | *-0.104 | *0.181 | *0.124 |
| reactIDR | **\*0.315** | **\*0.201** | **\*0.255** | *0.114 |

\* represents $p$ -value $< 10^{-5}$ after Bonferroni correction.

Moreover, we examined the accuracy of the reactivity scores from multiple HTS datasets for the same test set combined with SVM, a representative machine learning method. As a result, reactIDR demonstrated the highest mean prediction accuracy of reactIDR compared to other methods by merging reactivity features for all datasets (Figure 4). As reactIDR was also observed to generate the highest average accuracy with linear discriminant analysis instead of SVM (see also Supplementary Figures 12 and 13), these results suggests the robustness of reactIDR when used in combination with different HTS approaches, and the potential in integrative analyses of various HTS datasets regardless of classification method.

The reactIDR reactivity was further examined for *in silico* structure prediction using RNAfold (Lorenz *et al.*, 2011). We applied the previously developed pseudo-free energy method (Deigan *et al.*, 2009) to perform a hybrid structure prediction of computational and experimental structural analyses, in which the reactivity is accounted for in the log linear form as

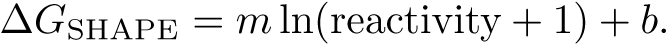

In Figure 6, PPV and sensitivity of the minimum free energy structure predicted by RNAfold with reactivity scores of reactIDR are shown, while progressively changing the parameters of the slope *m* and intercept *b*. To compare the PPV and sensitivity, the best PPV and sensitivity pair was selected to maximize the sum of pairs for *in silico* prediction of RNAfold with or without reactivity scores of reactIDR and PROBer. As a result, the reactivity scores of reactIDR with RNAfold were shown to perform better with the maximum PPV and sensitivity pair shown to be (0.290 and 0.373) while the best pair for *in silico* prediction was (0.214, 0.305) and that for PROBer was (0.249, 0.369) (Figure 5 and Supplementary Figure 14). The reactivity-guided structure prediction was also applied for the case in which only a partial reactivity information (0 % or no reactivity information, 10 %, 20 %, up to 100 % or all reactivity information) were provided and selection priority was given to top ranking reactivity scores. With the fixed parameters (*m* = 0.5 and *b* = 0.0) selected from the results presented in Figure 6, the Matthews correlation coefficient (MCC) of secondary structure prediction by using RNAfold were overall higher than the complete *in silico* structure prediction, even when only the top 10 % reactivity scores were provided as a guide (Supplementary Figure 15).

Taken together, the reactIDR reactivity scores were able to improve the prediction accuracy of integrative analyses as well as *in silico* structure prediction as conformational constraints. According to the assumptions in reactIDR, the reactivity scores correspond to the probability of the signals being true or false signals, which accounts for the reproducibility. Thus, it might be theoretically applicable to the cross-comparison of reactivity scores between the datasets obtained by different HTS methods. Hence, reactIDR may be useful for the optimization and validation of downstream analyses, such as *in silico* structure prediction, based on the filtering of unreliable information inferred by the unified model for multiple HTS methodologies.

### 3.3 Correlation between RNA reactivity and 3D accessibility

The agreement between reactivity and 2D accessibility was demonstrated for rRNAs, but the reactivity of RNA in a chemical probing reaction appears to be affected by the 3D conformation, as observed in some previous studies (Bindewald *et al.*, 2011). In particular, the presence of the sugar puckers of RNA structures in *C*2′- or *C*3′-end is considered to affect the modification efficiency of SHAPE reagent (Vicens *et al.*, 2007; Mlynsky and Bussi, 2017). To assess the influence of 3D accessibility on reactivity, we examined the correlation of the reactivity score with 2D and 3D accessibility. For 3D accessibility, ASA was computed for each nucleotide in the human ribosome structure using NACESS (Hubbard and Thornton, 1993).

In Figure 7(A), the relationship between ASA and reactivity score is presented, in which ASA is averaged with 500-nt sliding window estimates for the test set of rRNAs (selected in the previous analysis) for the *in vivo* icSHAPE dataset (also see Supplementary Figure 16). All analyzed methods showed a positive correlation between ASAs and reactivity score ranking (shown in Table 3. 2), with the exception of a partial region of ties appeared when using the BUMHMM or low reactive bases using the PROBer. Furthermore, reactIDR showed the highest correlation across the six scoring methods with a statistically significant *p*-value (*p <* 10^*-*^10^^ after Bonferroni multiple correction). This result was expected to be consistent with the high accuracy of stem/loop classification because 2D accessibility and 3D accessibility are also highly correlated (Supplementary Figure 17). On the other hand, the correlation between reactivity scores and 3D accessibility was lower for the *in vitro* dataset except for the moderate decline for reactIDR. We additionally investigated the correlation of reactivity with 2D accessibility by setting loop and stem structures to 1 and 0, showing a different tendency that the correlation between reactivity and 2D accessibility was positive in both of the *in vivo* and *in vitro* datasets across all reactivity scoring methods.

Using the HTS datasets, it is possible to confirm that the 3D structures *in vitro* are different from those observed *in vivo* and reference structures due to the lack of protein binding. The *in vivo* reactive sites (represented by red in Figure 7(B)) were observed to be located at the outer side of ribosomes compared to those obtained *in vitro*, and the regions with the highly-reactive nucleotides *in vivo* and *in vitro* rarely overlapped. Since the aim of reactIDR is to allow comparisons between different HTS datasets in a fair manner, the reactIDR reactivity can potentially be used for the quantification of conformational changes between the different conditions.

### 3.4 Further applications of reactIDR

In this study, reactIDR was shown to achieve an accuracy statistically comparable to that of PROBer, while its precision at the low-recall area was slightly lower than that of PROBer (Figure 4 and Supplementary Figure 10). The previous studies applied various methods of read normalization or trimming, adjusted to their own reactivity scores, as well as computing the reactivity scores. Therefore, an exhaustive search for an appropriate read preprocessing may increase the accuracy of structure prediction of our method.

Recent HTS experiments often are replicated and reactIDR requires HTS replication data for the reactivity computation. However, even if the target HTS dataset does not contain any replicates, a virtual replication by random division of datasets may be an option for reactIDR computation. In addition to the current scope of reactIDR in this study, novel HTS experiments can be incorporated as well, which have recently emerged based on the more complicated structural footprints, such as using mutational profiling (Siegfried *et al.*, 2014; Zubradt *et al.*, 2017) or base pair detection by RNA crosslink (Lu *et al.*, 2016; Sharma *et al.*, 2016). To increase the applicability of reactIDR for these HTS experiments, further extensions can be considered, such as the accounting for base type dependency, co-occurrence of base modification, or attenuation of read counts depending on the distance from the transcript ends. Although these applications would require a higher number of parameters to be able to consider more complicated contexts, the appropriate model can be learned using supervised learning with model selection methods.

## 4 Conclusions

We have developed a novel software, reactIDR, for the prediction of stem/loop regions from HTS analysis based on the reproducibility criterion. For the rRNA structure analyses, reactIDR achieved robust accuracy across different datasets. Moreover, the reactivity scores obtained with reactIDR enhanced the prediction accuracy by integrating multiple datasets and *in silico* structure analyses. In conclusion, reactIDR shows a great potential for the use in various HTS dataset analyses, enabling the progression of the current understanding of the RNA secondary structures.

## Acknowledgements

Computations were performed using the supercomputing facilities at the Human Genome Center, the University of Tokyo (http://supcom.hgc.jp/shirokane.html). We also thank Tony Kuo and Raissa Relator for critically reading the manuscript.

